# Genotype-environment interactions determine microbiota plasticity in *Nematostella vectensis*

**DOI:** 10.1101/2022.06.26.497683

**Authors:** Laura Baldassarre, Adam M. Reitzel, Sebastian Fraune

## Abstract

Most multicellular organisms harbor microbial colonizers that provide various benefits to their hosts. Although these microbial communities may be host species- or even genotype-specific, the associated bacterial communities can respond plastically to environmental changes. In this study, we estimated the relative contribution of environment and host genotype to bacterial community composition in *Nematostella vectensis*, an estuarine cnidarian. We isolated *N. vectensis* polyps from five different populations along a north-south gradient on the Atlantic coast of the United States and Canada at three different times of the year. While half of the polyps were immediately analyzed for their bacterial composition by 16S rRNA gene sequencing, the remaining polyps were cultured under laboratory conditions for one month. Bacterial community comparison analyses revealed that laboratory maintenance reduced bacterial diversity by fourfold, but maintained a population-specific bacterial colonization. Interestingly, the differences between bacterial communities correlated strongly with seasonal variations, especially with ambient water temperature. To decipher the contribution of both ambient temperature and host genotype to bacterial colonization, we generated 12 clonal lines from six different populations in order to maintain each genotype at three different temperatures for three months. The bacterial community composition of the same *N. vectensis* clone differed greatly between the three different temperatures, highlighting the contribution of ambient temperature to bacterial community composition. To a lesser extent, bacterial community composition varied between different genotypes under identical conditions, indicating the influence of host genotype. In addition, we identified a significant genotype x environment interaction determining microbiota plasticity in *N. vectensis*. From our results we can conclude that *N. vectensis*-associated bacterial communities respond plastically to changes in ambient temperature, with the association of different bacterial taxa depending in part on the host genotype. Future research will reveal how this genotype-specific microbiota plasticity affects the ability to cope with changing environmental conditions.

## Introduction

Most multicellular organisms live in association with microbial symbionts (McFall-Ngai et al., 2013; Zilber-Rosenberg & Rosenberg, 2008). It has been widely demonstrated that these symbionts provide various benefits for the survival and persistence of their hosts (Peixoto et al., 2017; Reshef et al., 2006; Rosado et al., 2019). The quality and quantity of associated microbial species is characteristic for host species (Baker, 2003; Fraune & Bosch, 2007; Kvennefors et al., 2010), genotype (Cahana & Iraqi, 2020; Glasl et al., 2019), biogeography (Linnenbrink et al., 2013; Mortzfeld et al., 2016; Terraneo et al., 2019), life stage (Baldassarre, Levy, et al., 2021; Damjanovic et al., 2020; Domin et al., 2018; Vijayan et al., 2019), diet (David et al., 2014; Leeming et al., 2019; Zarrinpar et al., 2014) and environmental conditions (Mortzfeld et al., 2016; Sehnal et al., 2021; Terraneo et al., 2019). Starting from these evidences, many studies demonstrated that the host plays an active role in shaping its symbiont microbiota (Augustin et al., 2017; Franzenburg et al., 2012; Fraune & Bosch, 2007; Groussin et al., 2017; Lee et al., 2016). In addition to the effects of the host and the environment, the interaction between these two factors is also discussed as a potential factor influencing the plasticity of the microbiota (Oyserman et al., 2021).

*Nematostella vectensis* is a small, burrowing estuarine sea anemone found in tidally restricted salt marsh pools. The distribution of this species extends over the Atlantic and Pacific coasts of North America, and the southeast coast of England (Reitzel et al., 2008) and its range encompasses large latitudinal variation in temperature and salinity (Sheader et al., 1997). *N. vectensis*’ s wide environmental tolerance and broad geographic distribution (Hand & Uhlinger, 1992; Reitzel et al., 2008), combined with the availability of a genome sequence (Putnam et al., 2007) make it an exceptional organism for exploring adaptations to variable environments. *N. vectensis* has separated sexes and it is able to reproduce both sexually through external fertilization (Darling et al., 2005; Hand & Uhlinger, 1992; Reitzel et al., 2007) and asexually through transverse fission (Reitzel et al., 2008). Although a free-swimming larval stage is present, this species is considered to have overall pretty limited dispersal abilities (Darling et al., 2004). Seasonal population fluctuations in density may lead to frequent bottlenecks, and when gene flow between subpopulations is restricted by physical barriers, such fluctuations could result in conspicuous genetic structuring of the metapopulation (Darling et al., 2004; Reitzel et al., 2010). Completely or largely clonal populations exist all through the distribution range of *N. vectensis* (Darling et al., 2004; Pearson et al., 2002; Reitzel et al., 2008), however microsatellite and SNP markers indicated an extensive intraspecific genetic diversity and genetic structuring between populations (Darling et al., 2009; Reitzel, Herrera, et al., 2013). Within a single estuary, *N. vectensis* occupies tidal streams that flush with each tide or, isolated still-water high-marsh pools, that can differ substantially in a set of ecological variables including temperature and salinity (Friedman et al., 2018; Smith & Able, 1994). Previous works showed that different *N. vectensis* genotypes from same natural pools, have significantly different tolerances to oxidative stress (Friedman et al., 2018) and that individuals from different field populations respond differently to same thermal conditions during lab culturing (Reitzel, Chu, et al., 2013).

An initial categorization of the *N. vectensis* microbiome has shown that individuals from different field pools of the North American Atlantic coast have significantly different microbiomes and that these differences follow a north-south gradient (Mortzfeld et al., 2016). The different ecological conditions that distinguish these pools from each other and the genetic structuring of *N. vectensis* populations led us to hypothesize that the microbiota is a subject to local selection. In particular, locally adapted host genotypes may associate with symbionts that provide advantages at the specific ecological conditions of each native pool.

In this study we analyzed the microbiota composition of polyps from different populations directly after sampling and after one month of laboratory maintenance. We first investigated which factors among ambient temperature, salinity, season and geographic location, contribute to microbiota differentiation. The results of these analyses show that the composition of the microbiota changes with both season and geographic location, and that these differences persist under laboratory conditions. Consistent with previous laboratory observations (Mortzfeld et al., 2016), our field data confirmed that temperature, not salinity, is correlating most with differences in bacterial community compositions. Starting from these evidences, we investigated the influence of ambient temperature on the microbiota plasticity of 12 individual genotypes derived from six different populations. We found that after three months of laboratory culture, temperature was the factor most driving microbiota differentiation, although differences according to genotype were also detectable. In addition, we demonstrated that microbiota plasticity in relation to temperature is genotype-specific, suggesting that microbiota plasticity is also influenced by interactions between genotype and temperature.

With this study, we have taken an important step toward understanding the contribution of both local environmental conditions and host genotype in shaping the microbiota. Furthermore, we have shown that, although the dynamics of the microbial community are plastic, each genotype is associated with a specific but limited plastic microbiota. These results suggest that local populations of the same species may have different abilities to adapt to environmental changes through microbiota-mediated plasticity.

## Materials and methods

### Animal sampling and culture

All experiments were carried out with polyps of *N. vectensis* (Stephenson 1935). Adult animals were collected from field populations of Nova Scotia (10/03/2016), Maine (11/03/2016, 02/06/2016, 11/09/2016), New Hampshire (11/03/2016, 02/06/2016, 11/09/2016), Massachusetts (12/03/2016, 03/06/2016, 13/09/2016), Maryland (long-term lab culture) and North Carolina (16/03/2016) by sieving them from loose sediments. Environmental parameters (air temperature, water temperature and salinity) were also recorded at the moment of sampling and used as metadata for further analysis (see **Table S1** for details). The animals from March sampling, were kept for one month under standard lab conditions and then the DNA was extracted. In the lab, the animals were kept under constant, artificial conditions, at 20°C, without substrate or light, in *N. vectensis* Medium (NM), which was adjusted to 16‰ salinity with Red Sea Salt^®^ and Millipore H_2_O. Polyps were fed 2 times a week with first instar nauplius larvae of *Artemia salina* as prey (Ocean Nutrition Micro *Artemia* Cysts 430 - 500 gr, Coralsands, Wiesbaden, Germany) and washed once a week with media pre-incubated at 20°C.

### Animal acclimation

Polyps from two different strains from each of the six wild populations, were placed separately, in three replicates, into 6 well plates and let acclimating for three months at three different temperatures (15, 20 and 25°C). After three months, the polyps were collected, frozen in liquid N and stored at -80°C before DNA extraction and 16S sequencing.

### DNA extraction

The specimens from the field were preserved in RNAlater until DNA extraction. For the samples from the field and after one month of lab culture, gDNA was extracted with the AllPrep DNA/RNA Mini Kit (Qiagen), as described in the manufacturer’s protocol. The animals from the experiment were washed two times with 2ml autoclaved MQ, instantly frozen in liquid N without liquid and stored at −80°C until extraction. The gDNA was extracted from whole animals with the DNeasy®Blood & Tissue Kit (Qiagen, Hilden, Germany), as described in the manufacturer’s protocol. Elution was done in 50μl and the eluate was stored at −80°C until sequencing. DNA concentration was measured by gel electrophoresis (5µl sample on 1.2% agarose) and by spectrophotometry through Nanodrop 3300 (Thermo Fisher Scientific).

### 16S rRNA sequencing

For each sample the hypervariable regions V1 and V2 of bacterial 16S rRNA genes were amplified. The forward primer (5’s- **AATGATACGGCGACCACCGAGATCTACAC** XXXXXXXX TATGGTAATTGT AGAGTTTGATCCTGGCTCAG-3’) and reverse primer (5’- **CAAGCAGAAGACGGCATACGAGAT** XXXXXXXX AGTCAGTCAGCC TGCTGCCTCCCGTAGGAGT -3’) contained the Illumina Adaptor (in bold) p5 (forward) and p7 (reverse). Both primers contain a unique 8 base index (index; designated as XXXXXXXX) to tag each PCR product. For the PCR, 100 ng of template DNA (measured with Qubit) were added to 25 µl PCR reactions, which were performed using Phusion® Hot Start II DNA Polymerase (Finnzymes, Espoo, Finland). All dilutions were carried out using certified DNA-free PCR water (JT Baker). PCRs were conducted with the following cycling conditions (98 °C—30 s, 30 × [98 °C—9s, 55 °C—60s, 72 °C—90s], 72 °C—10 min) and checked on a 1.5% agarose gel. The concentration of the amplicons was estimated using a Gel Doc TM XR+ System coupled with Image Lab TM Software (BioRad, Hercules, CA USA) with 3 µl of O’GeneRulerTM 100 bp Plus DNA Ladder (Thermo Fisher Scientific, Inc., Waltham, MA, USA) as the internal standard for band intensity measurement. The samples of individual gels were pooled into approximately equimolar subpools as indicated by band intensity and measured with the Qubit dsDNA br Assay Kit (Life Technologies GmbH, Darmstadt, Germany). Subpools were mixed in an equimolar fashion and stored at -20 °C until sequencing. Sequencing was performed on the Illumina MiSeq platform with v3 chemistry (Rausch et al., 2016). The raw data are deposited at the Sequence Read Archive (SRA) and available under the project PRJNA757926.

### Analyses of bacterial communities

The 16S rRNA gene amplicon sequence analysis was conducted through the QIIME 1.9.0 package (Caporaso et al., 2010). Sequences with at least 97% identity were grouped into OTUs and clustered against the QIIME reference sequence collection; any reads which did not hit the references, were clustered de novo. OTUs with less than 50 reads were removed from the dataset to avoid false positives which rely on the overall error rate of the sequencing method (Faith et al., 2013). Samples with less than 5500 sequences were also removed from the dataset, being considered as outliers. For the successive analysis the number of OTUs per sample was normalized to the lowest number of reads after filtering.

Alpha-diversity was calculated using the Chao1 metric implemented in QIIME. Beta diversity was calculated in QIIME according with the different beta-diversity metrics available (Binary-Pearson, Bray-Curtis, Pearson, Weighted-Unifrac and Unweighted-Unifrac). Statistical values of clustering were calculated using the nonparametric comparing categories methods Adonis and Anosim. MANOVA was performed through QIIME 2 (Bolyen et al., 2019).

Statistical tests were performed through JASP v0.16.2 (https://jasp-stats.org). Data were subjected to descriptive analysis, and normality and variance homogeneity tests as described herein. For univariate analyses, statistical differences were tested through non-parametric U-test due to the non-normal distribution of the data and the presence of significant outliers. For multivariate analyses, when normality, variance homogeneity and absence of significant outliers assumptions were met, statistical significance was tested through one-way ANOVA. When a significant difference between treatments was stated, Tukey’s post-hoc comparisons were performed in order to infer its direction and size effect. When at least one of the assumptions was violated, the non-parametric Kruskal-Wallis test was performed instead, followed by Dunn’s post-hoc comparisons.

Bacterial OTUs specifically associated with each genotype and each temperature were identified through LEfSe (http://huttenhower.sph.harvard.edu/galaxy) (Segata et al., 2011). LEfSe uses the non-parametric factorial Kruskal-Wallis sum-rank test to detect features with significant differential abundance, with respect to the biological conditions of interest; subsequently LEfSe uses Linear Discriminant Analysis (LDA) to estimate the effect size of each differentially abundant feature. Assuming that different genotypes from the same location may naturally share a number of symbionts, we only performed pairwise comparisons between genotypes from different locations. In addition to that, presence-absence calculations were performed directly on the OTU tables in order to detect bacterial OTUs that are unique for a specific genotype or AT.

## Results

### Laboratory maintenance results in loss of bacterial diversity associated with *N. vectensis* polyps

Genomic DNA samples from 168 *N. vectensis* polyps were submitted for 16S rRNA gene sequencing. While 53 samples were collected from 5 different populations (Nova Scotia, Maine, New Hampshire, Massachusetts and North Carolina) in March 2016, the sampling in Maine, New Hampshire and Massachusetts was repeated also in June and September (31 and 34 samples respectively). In addition, we maintained 50 polyps sampled in March, for one month under laboratory conditions before we extracted gDNA. Sequencing was successful for 156 samples. After picking OTUs represented by at least 50 reads, 6.431 different OTUs were detected, with 5.651 to 112.985 reads per sample.

Maintaining *N. vectensis* polyps for one month under laboratory conditions resulted in a major shift in the associated bacterial communities comparing to the bacterial communities of polyps directly sampled from the field (**Fig. 1A** and **Table 1**). While the bacterial variability between polyps significantly decreases during one month of laboratory culturing (**Fig. 1B**), the alpha-diversity decreases to around one quarter of the bacterial diversity observed in field sampled *N. vectensis* polyps (**Fig. 1C**).

**Table 1.** Statistical analysis determining the influence of animal laboratory maintenance on bacterial colonization. Statistical analyses were performed (methods ANOSIM and ADONIS, number of permutations = 999) on each of the pairwise comparison distance matrices generated.

| parameter | beta-diversity metric | Adonis |  | Anosim |  |
| --- | --- | --- | --- | --- | --- |
|  |  | *R*<sup>2</sup> | *p-value* | *R* | *p-value* |
| **Source** | Binary-Pearson | 0.219 | 0.001 | 0.857 | 0.001 |
|  | Bray-Curtis | 0.158 | 0.001 | 0.532 | 0.001 |
|  | Pearson | 0.139 | 0.001 | 0.278 | 0.001 |
|  | Weighted-Unifrac | 0.234 | 0.001 | 0.534 | 0.001 |
|  | Unweighted-Unifrac | 0.231 | 0.001 | 0.957 | 0.001 |

**Figure 1.**
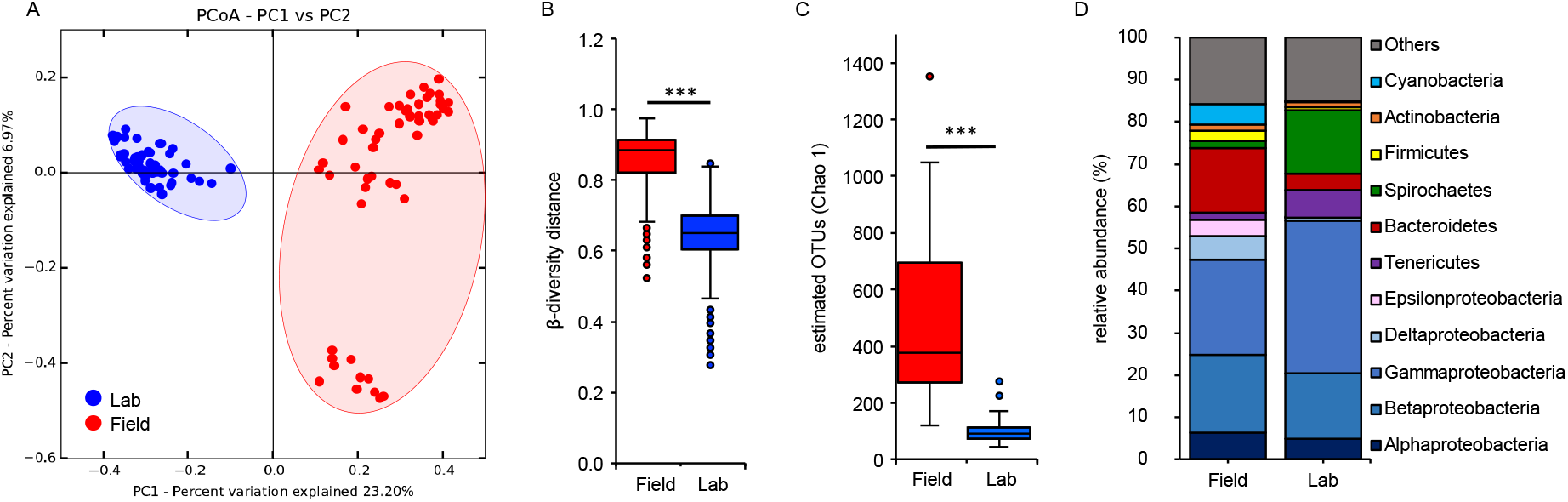
Laboratory maintenance reduced bacterial diversity associated with *N. vectensis* polyps. (A) PCoA (based on Binary-Pearson metric, sampling depth = 5500) illustrating similarity of bacterial communities based on sample source; (B) beta-diversity distance box-plots of the field and lab samples; (C) alpha-diversity (Chao1) comparisons between field and lab samples (max rarefaction depth = 5500, num. steps = 10). Differences in B and C were tested through Mann-Whitney U-test (*** = p ≤ 0.001). (D) relative abundance of main bacterial groups among the two different samples sources.

The loss of bacterial diversity in laboratory-maintained polyps became also evident by comparing the major bacterial groups (**Fig. 1D**). While Cyanobacteria and Epsilonproteobacteria disappeared and Firmicutes, Deltaproteobacteria, and Bacteroidetes decreased in relative abundance in laboratory-maintained animals, Spirochaetes and Tenericutes increased in relative abundance (**Fig. 1D**).

To determine whether bacterial communities from polyps collected from different locations reveal a biogeographic signal, and to test whether this signal is preserved in polyps maintained in the laboratory, we analyzed the two data sets, field and laboratory samples, separately.

### Microbial diversity in the field correlates with host biogeography and environmental factors

Analyzing the bacterial communities associated with the field sampled Nematostella polyps, principal coordinates analysis (PCoA) revealed a clear clustering of the associated bacterial community by provenance location (**Fig. 2A, B, Table 2**).

**Table 2.** Statistical analysis determining the influence of geographic location on bacterial colonization in field samples. Statistical analyses were performed (methods ANOSIM and ADONIS) on each of the pairwise comparison distance matrices generated (Number of permutations = 999).

| parameter | beta-diversity metric | Adonis |  | Anosim |  |
| --- | --- | --- | --- | --- | --- |
|  |  | R2 | p-value | R | p-value |
| Geographic location | Binary-Pearson | 0.322 | 0.001 | 0.862 | 0.001 |
|  | Bray-Curtis | 0.360 | 0.001 | 0.680 | 0.001 |
|  | Pearson | 0.410 | 0.001 | 0.445 | 0.001 |
|  | Weighted-Unifrac | 0.462 | 0.001 | 0.591 | 0.001 |
|  | Unweighted-Unifrac | 0.222 | 0.001 | 0.578 | 0.001 |

**Figure 2.**
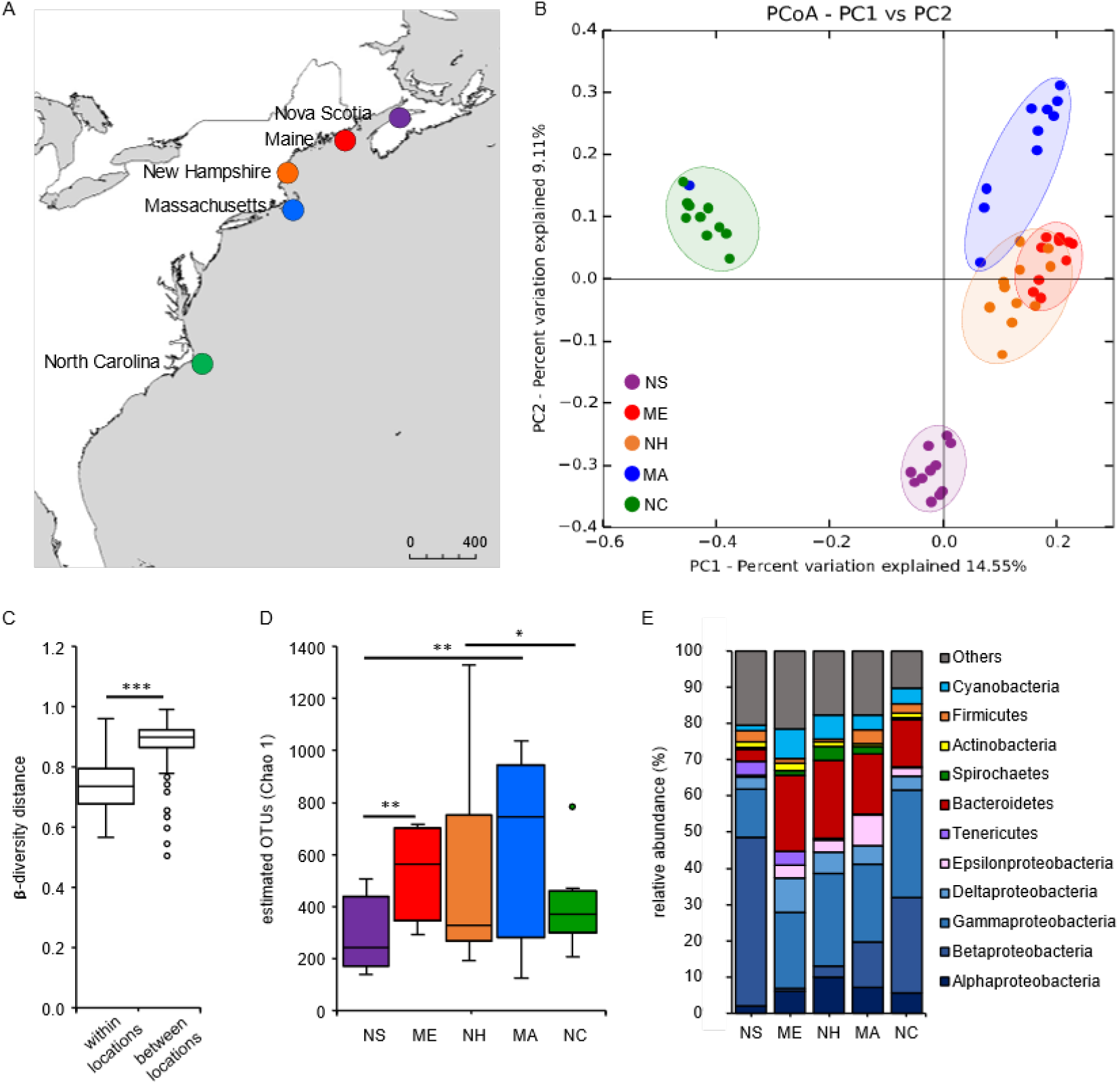
Natural *N. vectensis* populations are associated with specific microbiota. (A) Sampling sites map; (B) PCoA (based on Binary-Pearson metric, sampling depth = 5500) illustrating similarity of bacterial communities based on geographic location; (C) beta-diversity distance box-plots of the field samples within and between geographic locations, differences were tested through Mann-Whitney U-test (*** = p ≤ 0.001); (D) alpha-diversity (Chao1) comparisons between geographic locations (max rarefaction depth = 5500, num. steps = 10), differences were tested through Kruskal-Wallis test followed by Dunn’s post-hoc comparisons (H = 10.66, * = p ≤ 0.05, ** = p ≤ 0.01); (E) relative abundance of main bacterial groups among different geographic locations. NS (Nova Scotia), ME (Maine), NH (New Hampshire), MA (Massachusetts), NC (North Carolina).

Based on the different beta diversity measures, geographic location explained between 44% and 86% of the bacterial variability (**Table 2**). The beta-diversity distance between samples within the same location was significantly lower than that between the different locations, stressing the clustering of the samples sharing the same provenance (**Fig. 2C**).

We next investigated the influence of geographic distance, water temperature and water salinity on a continuous scale by applying Mantel tests to each of the five measures of beta-diversity (**Table 3**). Mantel tests revealed that the geographic distance is the main factor impacting beta-diversity, explaining approximately 20–60% of the variation in four out of five beta-diversity measures (**Table 3**). Only the Pearson metric revealed no significant correlation between geographic distance and bacterial composition. While both environmental factors temperature and salinity also correlated significantly with bacterial diversity, water temperature explained a higher proportion of bacterial diversity (**Table 3**).

**Table 3.** Statistical analysis determining the influence of geographic distance, field temperature and salinity on bacterial colonization. Mantel tests were performed between the three different parameters distance matrices and the beta-diversity matrices generated. (Number of permutations = 999).

| parameter | beta-diversity metric | Mantel test |  |
| --- | --- | --- | --- |
|  |  | r | p-value |
| Geographic distance | Binary-Pearson | 0.603 | 0.001 |
|  | Bray-Curtis | 0.199 | 0.002 |
|  | Pearson | -0.045 | 0.473 |
|  | Weighted-Unifrac | 0.270 | 0.001 |
|  | Unweighted-Unifrac | 0.362 | 0.001 |
| Temperature | Binary-Pearson | 0.592 | 0.001 |
|  | Bray-Curtis | 0.210 | 0.001 |
|  | Pearson | -0.024 | 0.638 |
|  | Weighted-Unifrac | 0.264 | 0.001 |
|  | Unweighted-Unifrac | 0.381 | 0.001 |
| Salinity | Binary-Pearson | 0.307 | 0.001 |
|  | Bray-Curtis | 0.381 | 0.001 |
|  | Pearson | 0.297 | 0.001 |
|  | Weighted-Unifrac | 0.162 | 0.002 |
|  | Unweighted-Unifrac | 0.149 | 0.005 |

In addition, alpha-diversity showed also a biogeographic signal. Polyps from the extreme northern and southern locations (Nova Scotia and North Carolina) had lower bacterial alpha-diversity than polyps from central locations (**Fig. 2D**). By looking at the principal bacterial groups in the field samples, a north-south pattern was evident regarding the Betaproteobacteria that increased in relative abundance moving from Maine through North Carolina, while Tenericutes and the bacteria comprised in the category “Others”, decreased in abundance moving in the same direction. The samples from Nova Scotia showed a different trend, with the Betaproteobacteria reaching the highest overall abundance while the Epsilonproteobacteria, Bacteroidetes and Cyanobacteria the lowest (**Fig. 2E**).

For the locations in which the samplings have been repeated at three different seasonal time points (Maine, New Hampshire and Massachusetts), we investigated the differences in the microbiota composition according to sampling month (March, June and September). A clustering of the samples with sampling time point was significant (**Fig. 3A**), contributing up to 39% of the total difference (**Table 4**). Interestingly, the samples from June clustered in between those from March and September (**Fig. 3A**), and showed a significant higher alpha-diversity than the other two sampling time points, suggesting a gradual shift of associated bacteria along seasons (**Fig. 3B**). The bacteria grouped under the category “Others” increased in abundance moving from March to September in all the three locations (Maine, New Hampshire and Massachusetts). Overall, the Bacteroidetes and Deltaproteobacteria were more abundant in March samples, while Cyanobacteria were more abundant and Gammaproteobacteria less abundant in the samples from June, respectively. Interestingly, Betaproteobacteria were detectable only in the samples from March and Tenericutes almost disappeared in September. The Firmicutes and Actinobacteria reached their maximum abundances in September, while Spirochaetes in June, together with Epsilonproteobacteria (**Fig. 3C**).

**Table 4.** Statistical analysis determining the influence of season on bacterial colonization. Statistical analyses were performed (methods ANOSIM and ADONIS, number of permutations = 999) on each of the pairwise comparison distance matrices generated.

| parameter | beta-diversity metric | Adonis |  | Anosim |  |
| --- | --- | --- | --- | --- | --- |
|  |  | *R*<sup>2</sup> | *p*-value | *R* | *p*-value |
| Season | Binary-Pearson | 0.120 | 0.001 | 0.350 | 0.001 |
|  | Bray-Curtis | 0.160 | 0.001 | 0.390 | 0.001 |
|  | Pearson | 0.274 | 0.001 | 0.389 | 0.001 |
|  | Weighted-Unifrac | 0.220 | 0.001 | 0.380 | 0.001 |
|  | Unweighted-Unifrac | 0.080 | 0.001 | 0.220 | 0.001 |

**Figure 3.**
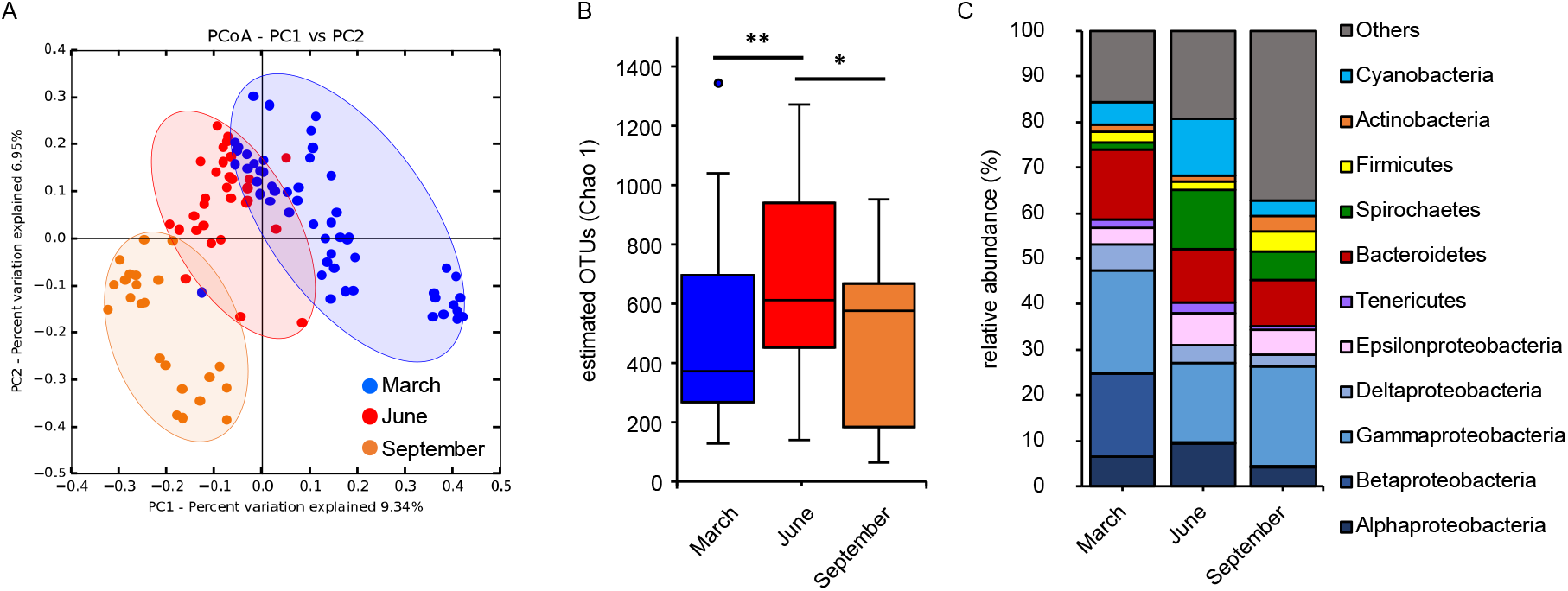
Natural microbiota in N. vectensis vary according to season. (A) PCoA (based on Binary-Pearson metric, sampling depth = 5500) illustrating similarity of bacterial communities based on sampling month; (B) alpha-diversity (Chao1) comparisons between sampling months (max rarefaction depth = 5500, num. steps = 10), differences were tested through Kruskal-Wallis test followed by Dunn’s post-hoc comparisons (H = 6.991, * = p = 0.015, ** = p = 0.008); (C) relative abundance of main bacterial groups among different sampling months.

### *N. vectensis* polyps cultured in the laboratory maintain population-specific microbiota

To test whether the biogeographic signal of the bacterial communities associated with polyps is maintained under laboratory conditions, we analyzed the laboratory samples separately (**Fig. 4**). A clear clustering of the samples according with the provenance location was still present and become even more evident after one month under laboratory conditions (**Fig. 4A** and **B**). All the ANOVA comparisons performed and the Mantel tests were highly significant (p < 0.001) (**Table 5**), and showed that the provenance geographic location explained between 50 and 65% of the beta-diversity difference for the lab samples, proving that the population-specific bacterial fingerprints were maintained (**Table 5**). The beta-diversity distance between samples originating from the same location was significantly lower than that between the different locations, stressing the clustering of the samples sharing the same provenance (**Fig. 4B**). For the lab samples, the alpha-diversity was also the highest in the samples from the intermediate locations (**Fig. 4C**). Animals from the extreme locations (Nova Scotia and North Carolina) where colonized by the highest abundances of Betaproteobacteria and Tenericutes, while those from the central latitudes were associated mainly with Gammaproteobacteria and Spirochaetes (**Fig. 4D**).

**Table 5.** Statistical analysis determining the influence of geographic distance and geographic location on bacterial colonization in laboratory-maintained populations. Statistical analyses were performed (methods ANOSIM and ADONIS) on each of the pairwise comparison distance matrices generated according with provenance geographic location. Mantel test was performed between the geographic location distance matrix and the different beta-diversity matrices. (Number of permutations = 999).

| parameter | beta-diversity metric | Adonis |  | Anosim |  | Mantel test |  |
| --- | --- | --- | --- | --- | --- | --- | --- |
|  |  | R2 | p-value | R | p-value | r | p-value |
| Geographic location/distance | Binary-Pearson | 0.283 | 0.001 | 0.656 | 0.001 | 0.287 | 0.001 |
|  | Bray-Curtis | 0.443 | 0.001 | 0.608 | 0.001 | 0.304 | 0.001 |
|  | Pearson | 0.632 | 0.001 | 0.609 | 0.001 | 0.424 | 0.001 |
|  | Weighted-Unifrac | 0.427 | 0.001 | 0.500 | 0.001 | 0.388 | 0.001 |
|  | Unweighted-Unifrac | 0.283 | 0.001 | 0.641 | 0.001 | 0.383 | 0.001 |

**Figure 4.**
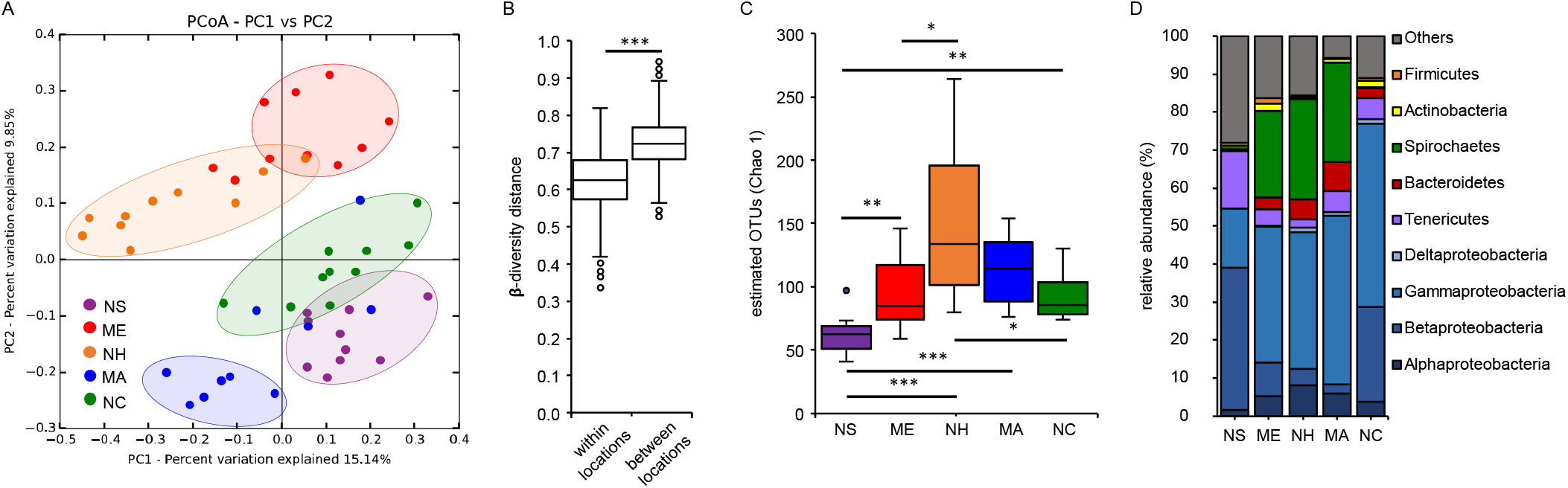
Population-specific microbiota are maintained under laboratory conditions. (A) PCoA (based on Binary-Pearson metric, sampling depth = 5500) illustrating similarity of bacterial communities based on geographic population; (B) beta-diversity distance box-plots of the lab samples within and between geographic locations, differences were tested through Mann-Whitney U-test (*** = p ≤ 0.001); (C) alpha-diversity (Chao1) comparisons between geographic locations (max rarefaction depth = 5500, num. steps = 10); (D) relative abundance of main bacterial groups among different geographic locations. Differences were tested through Kruskal-Wallis test followed by Dunn’s post-hoc comparisons (H = 25.2, * = p ≤ 0.05, ** = p ≤ 0.01, *** = p ≤ 0.001). NS (Nova Scotia), ME (Maine), NH (New Hampshire), MA (Massachusetts), NC (North Carolina).

### Under different temperatures, *N. vectensis* maintains genotype-specific microbiota

The variation of bacterial communities associated with *N. vectensis* polyps in the field correlated mostly with ambient water temperature (**Table 3**). Based on these findings, we aimed to measure experimentally the contribution of temperature and host genotype and their interaction on the microbiota composition. We selected in total 12 genotypes originating form 6 different geographic locations (2 genotypes/location) (**Fig. 5A**). To be able to maintain each genotype at different ambient temperatures, we clonally propagated the polyps to reach at least 9 clones/genotype. Subsequently, we maintained each genotype at three different temperatures (15, 20 and 25°C, n=3) for three months (**Fig. 5A**). Nine polyps out of 108 didn’t survive the treatment. Interestingly, culturing at high temperature (25°C) resulted in higher mortality in animals from Nova Scotia, New Hampshire, and Massachusetts, while animals from Maine had the highest mortality at low temperatures (15 and 20°C) (**Fig. S1**).

After three months of culturing at different temperatures, gDNA from 99 polyps were submitted for 16S rRNA gene sequencing. After picking up OTUs represented by at least 50 reads, 1.093 different OTUs were detected, with the number of reads per sample ranging between a maximum of 75.806 and a minimum of 3.156. After setting the minimum number of reads/sample at 11.700, 92 samples remained for the successive analyses.

PcoA revealed that ambient temperature explained most of the detected bacterial diversity associated with the polyps (between 38% and 67% diversity explained) (**Fig. 5B, Table 6)**, and that the 20°C samples had higher beta-diversity variability compared to polyps maintained at the two extreme temperatures (**Fig. 5C**). While principal component 1 (PC1) mostly separates samples according to the ambient temperature (**Fig. 5B**), PC2 mostly explains variations within the different genotypes (**Fig. 5E**). The ANOSIM results indicated that host genotype contributed between 8% and 20% to the total bacterial diversity observed (**Table 6**). While animals acclimated at 20°C showed a significant higher alpha-diversity than those acclimated at 15 and 25°C (**Fig. 5D**), no effects from the host genotypes on the alpha-diversity analysis were evident (**Fig. 5F**).

**Table 6.** Statistical analysis determining the influence of host genotype and temperature on bacterial colonization in experimental animals. Statistical analyses were performed (methods ANOSIM and ADONIS, number of permutations = 999) on each of the pairwise comparison distance matrices generated.

| parameter | beta-diversity metric | Adonis |  | Anosim |  |
| --- | --- | --- | --- | --- | --- |
|  |  | *R*<sup>2</sup> | *p*-value | *R* | *p*-value |
| Temperature | Binary-Pearson | 0.188 | 0.001 | 0.671 | 0.001 |
|  | Bray-Curtis | 0.193 | 0.001 | 0.500 | 0.001 |
|  | Pearson | 0.212 | 0.001 | 0.385 | 0.001 |
|  | Weighted-Unifrac | 0.436 | 0.001 | 0.545 | 0.001 |
|  | Unweighted-Unifrac | 0.165 | 0.001 | 0.453 | 0.001 |
| Genotype | Binary-Pearson | 0.202 | 0.001 | 0.167 | 0.001 |
|  | Bray-Curtis | 0.230 | 0.001 | 0.188 | 0.001 |
|  | Pearson | 0.277 | 0.001 | 0.204 | 0.001 |
|  | Weighted-Unifrac | 0.160 | 0.137 | 0.022 | 0.232 |
|  | Unweighted-Unifrac | 0.170 | 0.003 | 0.080 | 0.014 |

In order to detect genotype-specific bacterial adjustments to temperature variation, we performed MANOVA analyses confirming that in addition to the individual effects of temperature and genotype, also genotype x temperature interactions significantly influenced microbial plasticity (**Table 7**). Plotting the average PC 2 eigenvalues of each genotype at the three different ambient temperatures **(Fig. 5G)**, revealed that the microbial plasticity differed between the 12 different genotypes. Interestingly, the adjustments in bacterial diversity within the 12 genotypes can be divided in two main patterns (**Fig. S4 A and B**) suggesting different metaorganism strategies to cope with environmental changes.

**Figure 1.**
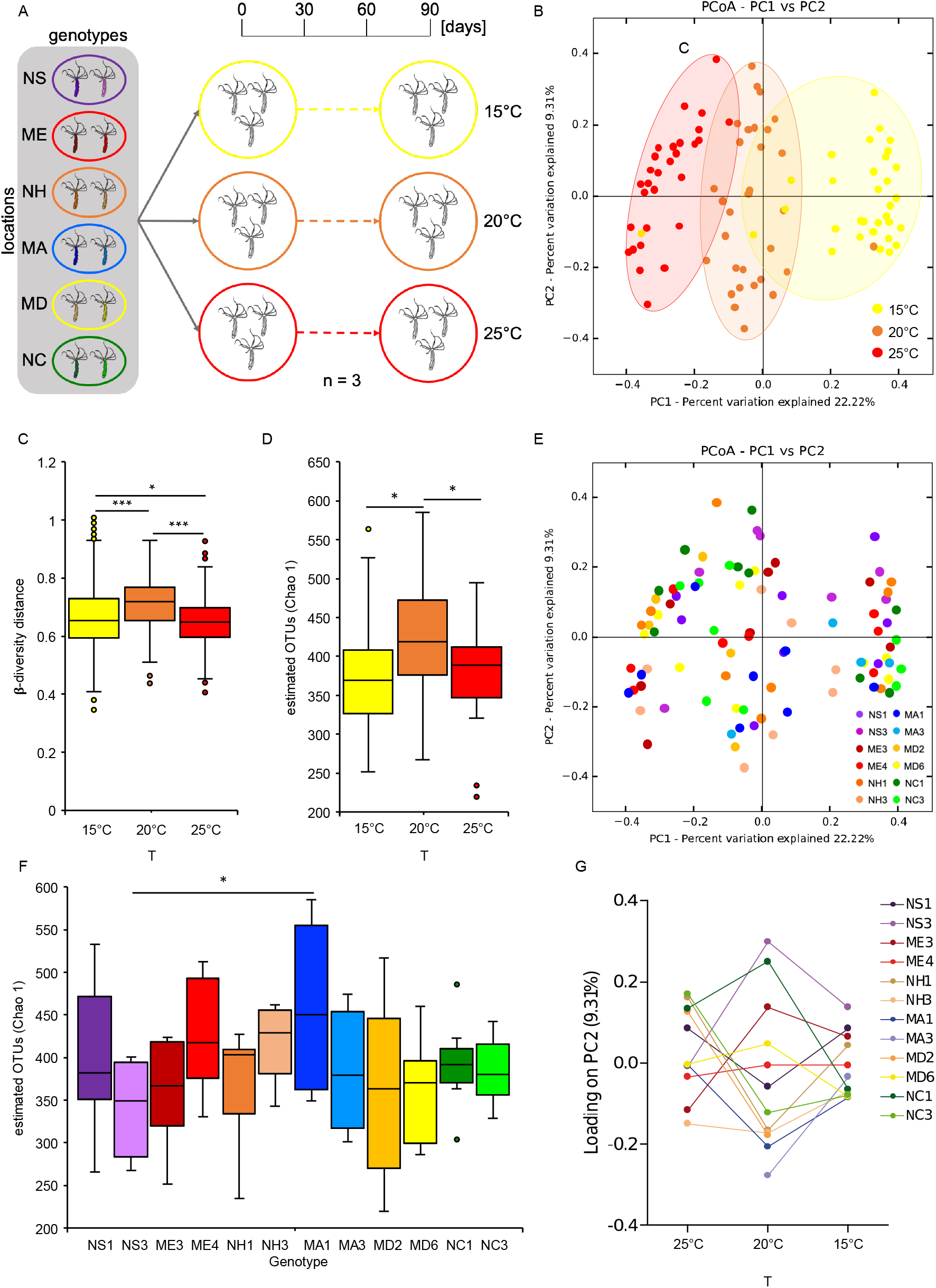
Influence of host genotype and temperature on bacterial colonization. (A) Experimental design, 2 genotypes for each geographic location were kept in 3 replicates at 3 different temperatures for 3 months; (B) PcoA (based on Binary-Pearson metric, sampling depth = 11700) illustrating similarity of bacterial communities based on ambient temperature; (C) alpha-diversity (Chao1) comparisons between temperatures (max rarefaction depth = 11700, num. steps = 10), differences were tested through one-way ANOVA followed by Tukey’s post-hoc comparisons (F = 4.313, * = p ≤ 0.05); (D) beta-diversity distance box-plots between different temperatures, differences were tested through Kruskal-Wallis test followed by Dunn’s post-hoc comparisons (H = 136.76, * = p = 0.024, *** = p ≤ 0.001); (E) PCoA (based on Binary-Pearson metric, sampling depth = 11700) illustrating similarity of bacterial communities based on host genotype; (F) alpha-diversity (Chao1) comparisons between polyp genotypes (max rarefaction depth = 11700, num. steps = 10), differences were tested through one-way ANOVA followed by Tukey’s post-hoc comparisons (F = 2.126, * = p = 0.027); (G) Reaction norms plotting average principal component 2 eigenvalues for each of the twelve genotypes at each temperature. NS (Nova Scotia), ME (Maine), NH (New Hampshire), MA (Massachusetts), MD (Maryland), NC (North Carolina), numbers near the location abbreviations indicate the different genotypes.

**Table 7.** Statistical analysis determining the influence of host genotype x temperature interaction on bacterial colonization in experimental animals. MANOVA test was performed (number of permutations = 999) on each of the beta-diversity distance matrices generated.

| parameter | beta-diversity metric | MANOVA |  |
| --- | --- | --- | --- |
|  |  | *R*<sup>2</sup> | *P* value |
| Temperature * Genotype | Binary-Pearson | 0.239 | 0.001 |
|  | Bray-Curtis | 0.221 | 0.001 |
|  | Pearson | 0.212 | 0.001 |
|  | Weighted-Unifrac | 0.167 | 0.008 |
|  | Unweighted-Unifrac | 0.234 | 0.001 |
| Temperature | Binary-Pearson | 0.223 | 0.001 |
|  | Bray-Curtis | 0.230 | 0.001 |
|  | Pearson | 0.253 | 0.001 |
|  | Weighted-Unifrac | 0.451 | 0.001 |
|  | Unweighted-Unifrac | 0.197 | 0.001 |
| Genotype | Binary-Pearson | 0.187 | 0.001 |
|  | Bray-Curtis | 0.217 | 0.001 |
|  | Pearson | 0.265 | 0.001 |
|  | Weighted-Unifrac | 0.112 | 0.009 |
|  | Unweighted-Unifrac | 0.153 | 0.001 |

In a further step, we aimed to detect indicator taxa specifically associated with ambient temperature and genotypes **(Fig. 6 and Table S2 and S3)**. Through LEfSe we were able to detect indicator OTUs that are overrepresented in each sample category in comparison with all the others. We observed that extreme ambient temperatures and genotypes from middle locations showed higher numbers of unique associated OTUs (**Fig. 6A and B**). Interestingly, calculating the relative abundance of indicator OTUs (**Fig. 6C and D**) revealed that around 37% and 27% of bacterial abundance at 15°C and 25°C respectively, were represented by temperature specific OTUs. In contrast genotype-specific OTUs represented on average 5-10% of the bacterial total abundance, while the two genotypes isolated from MD (the only long-term lab culture) were mainly associated with one genotype-specific OTU (belonging to the Spirochetes) representing around 30-40% of the total bacterial abundance (**Fig. 6D**). Interestingly, genotypes isolated from the same location show high similarities in terms of specific OTUs and their relative abundances (**Fig. 6C and D**), suggesting that genotypes from the same locations might be close realtives.

**Figure 2.**
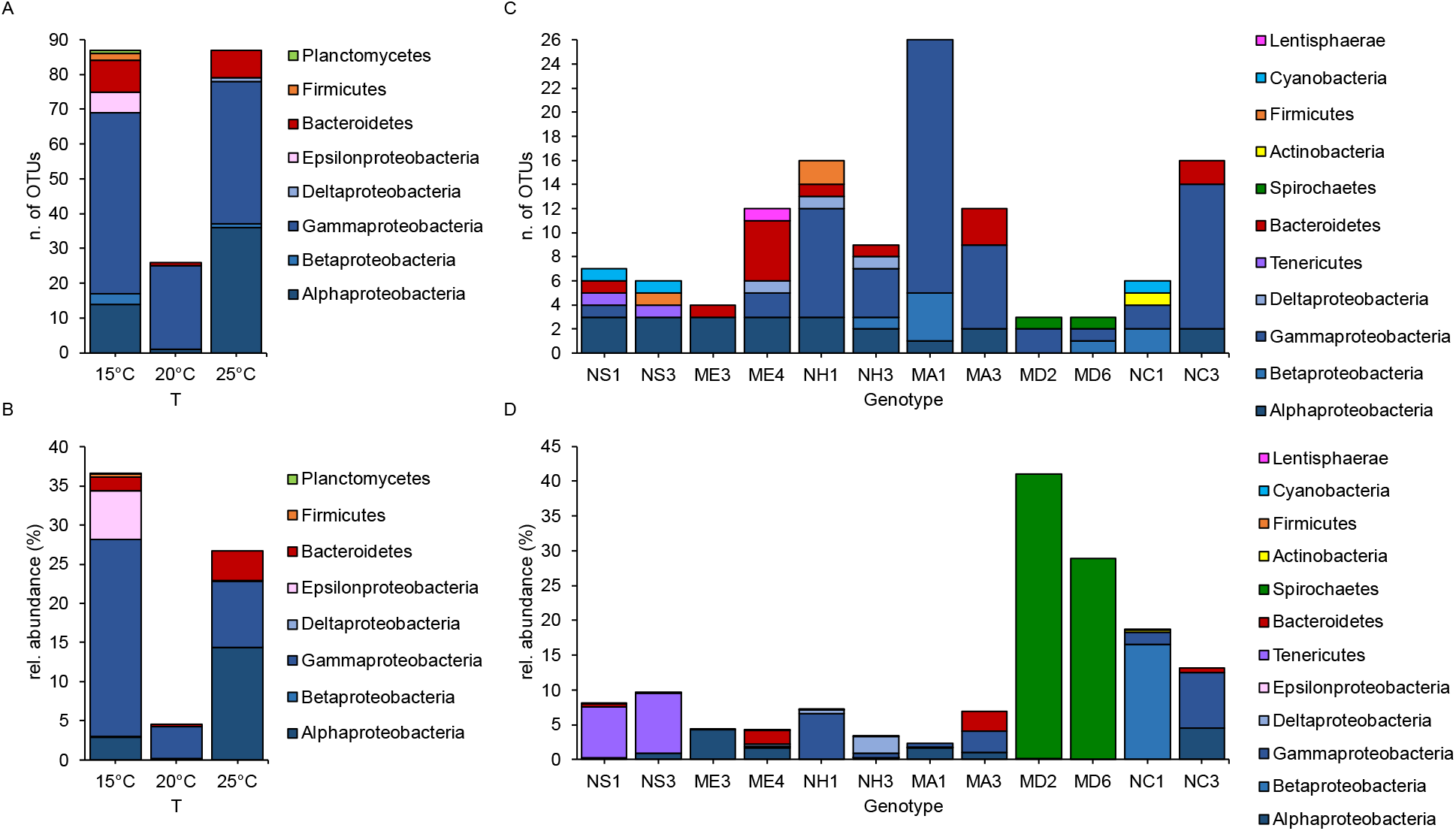
Bacterial OTUs representative of host genotype and acclimation temperature. Number of bacterial OTUs overrepresented at each temperature (A and B) and in each genotype (C and D) compared to the others, divided by major groups. Absolute OTU number (A and C), relative OTUs abundances on the total number of reads (B and D).

## Discussion

### Environmental factors can explain most but not all variability of *N. vectensis* associated microbiota

To estimate the contributions of both environmental factors and genotype to the bacterial diversity associated with *N. vectensis*, we started with a huge sampling effort to collect individuals of *N. vectensis* from all the main populations of the US Atlantic coast along a north south gradient of more than 1500 km and correlated the microbial composition data to the environmental factors’ temperature and salinity. In addition, we sampled individuals from three populations also in three different seasons. Our results showed that temperature and salinity, although explaining a similar percentage of the observed variability, could not explain all of the observed bacterial variation. In addition, we showed that the associated microbial community changes gradually along a temporal pattern during the year. Previous studies in corals have also shown that associated bacterial communities change depending on the season (Cai et al., 2018; Li et al., 2014; Rubio-Portillo et al., 2016; Sharp et al., 2017), e.g., due to changes in dissolved oxygen concentrations and rainfall (Li et al., 2014). In addition, seasonal changes in host physiology associated with winter quiescence, may drive microbiota diversity (Sharp et al., 2017). Besides these cues, natural seasonal fluctuations in bacterial communities can also impact the availability of certain symbiotic species (La Rivière et al., 2013).

### Maintenance in the laboratory reduces bacterial diversity but preserves population-specific bacterial signatures

After sampling polyps from the field, we additionally kept individuals of *N. vectensis* from each population under constant laboratory conditions for one month and compared these samples to those sampled directly from the field in terms of microbial diversity. In accordance with what was previously found from studies on lab-mice (Bowerman et al., 2021) and insects (Ibarra-Juarez et al., 2018; Morrow et al., 2015; Staubach et al., 2013), laboratory-reared *N. vectensis* individuals host a significantly lower bacterial diversity than in the wild. Interestingly, the homogenous lab environment did not eliminate the original differences in bacterial colonization observed in the animals directly samples from the field. Surprisingly, the population specific signature became even more evident in the laboratory-maintained animals. These results indicate that the bacterial diversity loss mainly affects bacteria that are not responsible for the population-specific signature. Therefore, bacteria that are lost under laboratory condition most likely are loosely associated environmental bacteria, food bacteria or might stem from taxa that are only transiently associated with the host. Bacteria that are persisting during laboratory maintenance most likely represent bacteria that are functionally associated with *N. vectensis* and might have co-evolved with its host.

### Genotype x environment interactions shape microbiota plasticity of *N. vectensis*

For several animal and plant species it has been observed that associated microbial community dissimilarities increase with geographical distance (Dunphy et al., 2019). Host selection, environmental filtering, microbial dispersal limitation and microbial species interactions have all been suggested as key drivers of host-microbial composition in space and time (Costello et al., 2012). Also a previous study in *N. vectensis* evidenced that individuals from different populations harbor distinct microbiota (Mortzfeld et al., 2016).

In order to disentangle the contribution of the host, the environment, and their interaction on the microbiota composition in *N. vectensis*, we selected 12 genotypes from six different field populations and kept clones of each genotype for three months under different temperatures. We found bacterial taxa that are associated with both specific genotypes and specific temperature conditions. These results suggest that both intrinsic and extrinsic factors shape the host-associated microbiota, although environmental conditions appear to have a stronger influence. In contrast to previous observations in corals (Brener-Raffalli et al., 2018; Glasl et al., 2019), where host genotype had a greater impact on microbiota composition than environmental conditions, in our study we observed that environmental conditions (in this case, temperature), tend to even the microbiota of different genotypes. Similar results were shown in fire coral clones, where both host genotype and reef habitat contributed to bacterial community variabilities (Dubé et al., 2021). Genomic function predictions suggested that environmentally determined taxa lead to functional restructuring of the microbial metabolic network, whereas bacteria determined by host genotype are functionally redundant (Dubé et al., 2021). As previously suggested (Damjanovic et al., 2019), these observations confirm that both environmental and host factors are drivers of associated microbial community composition and that different genotype x environment combinations can create unique microhabitats suitable for different microbial species with different functions.

One mechanism by which host selection can occur is through innate immunity, e. g. the secretion of antibiotic compounds via the mucus layer that target non-beneficial or pathogenic microbes (Augustin et al., 2017; Franzenburg et al., 2012; Fraune & Bosch, 2007; Ritchie, 2006). Our results suggest that *N. vectensis* also plays an active role in shaping its symbiotic microbiota in response to environmental variability and that these mechanisms depend on genotypic differences and local adaptation.

### Microbial plasticity is promoting animal adaptations

Differences in prokaryotic community composition in different environments have been documented in many other marine invertebrates and are considered to reflect local acclimation (A Hernandez-Agreda et al., 2016; Glasl et al., 2019; Goldsmith et al., 2018; van Oppen et al., 2018). We have recently shown that the restructuring of microbial communities due to temperature acclimation is an important mechanism of host plasticity and adaptation in *N. vectensis* (Baldassarre, Ying, et al., 2021). The higher thermal tolerance of animals acclimated to high temperature could be transferred to non-acclimated animals through microbiota transplantation (Baldassarre, Ying, et al., 2021). In our study, high temperature conditions were particularly challenging for some genotypes native to north habitats, where they experience colder climate. Whether this is the result of local adaptation of the host to colder temperatures or the symbiotic microbiota, needs to be clarified. We also observed that the bacterial species richness increases in intermediate latitudes, seasons and temperature, while it decreases at the extremes, suggesting a dynamic and continuous remodeling of the microbiota composition along environmental conditions gradients.

Evidence from reciprocal transplantation experiments in corals followed by short-term heat stress suggests also that coral-associated bacterial communities are linked to variation in host heat tolerance (Ziegler et al., 2017) and that associated bacterial community structure responds to environmental change in a host species-specific manner (Ziegler et al., 2019). Therefore, we hypothesize that host organisms may evolve faster than on their own due to plastic changes in their microbiota. Rapidly dividing microbes are predicted to undergo adaptive evolution within weeks to months. Adaptation of the microbiota can occur via changes in absolute abundancies of specific members, acquisition of novel genes, mutation and/or horizontal gene transfer (Baldassarre, Ying, et al., 2021; Bang et al., 2018; Bay & Palumbi, 2015; Edwards, 2020; van Oppen et al., 2018). Here, we provide evidence for genotype-specific microbial plasticity, leading to genotype-specific restructuring of the microbial network in response to environmental stimuli. Together these results may indicate that the genotype-specific bacterial colonization reflects local adaptation. In particular, genotypes adapted to highly variable environments might favor flexibility over fidelity regarding the associated microbiota composition; conversely, under more stable conditions less dynamic and more strict association might be advantageous (Voolstra & Ziegler, 2020).

## Acknowledgments

This work was supported by the Human Frontier Science Program (Young Investigators’ Grant RGY0079/2016 and the DFG CRC grant 1182 “Origin and Function of Metaorganisms” (Project B1). We thank Katja Cloppenborg-Schmidt (CRC 1182 project Z3) for preparing the 16S rRNA gene library and the Institute of Clinical Molecular Biology in Kiel for providing sequencing services.

## Conflict of Interest statement

The authors declare no conflict of interest.

Supporting information file is available online.

## Supplementary Material

**Table S1.** Metadata and environmental data at sampling time points.

| Location | Latitude | Longitude | Collection Date | Temperature (C°) |  |  | Salinity (‰) |
| --- | --- | --- | --- | --- | --- | --- | --- |
|  |  |  |  | Avg | Max | SD |  |
| Crescent Beach, Nova Scotia | 45154713 | -64371314 | 10/03/16 | -2.20* | 1.76** | // | 28 |
| Saco, Maine | 43560682 | -70271087 | 11/03/16 | 1.95 | 5.34 | 1.93 | 24 |
| Saco, Maine | 43560682 | -70271087 | 02/06/16 | 24.63 | 27.67 | 1.98 | 23 |
| Saco, Maine | 43560682 | -70271087 | 11/09/16 | 22.48 | 25.42 | 1.92 | 28 |
| Odiorne, New Hampshire | 42964569 | -70764588 | 11/03/16 | 10.15* | 13.42 | // | 35 |
| Odiorne, New Hampshire | 42964569 | -70764588 | 02/06/16 | 22.4* | 22.4 | // | 38 |
| Odiorne, New Hampshire | 42964569 | -70764588 | 11/09/16 | 28.12* | 29.76 | // | 35 |
| Sippewissett, Massachusetts | 41577021 | -70640749 | 12/03/16 | 12.23 | 19.47 | 4.8 | 22 |
| Sippewissett, Massachusetts | 41577021 | -70640749 | 03/06/16 | 21.71 | 26.2 | 2.81 | 24 |
| Sippewissett, Massachusetts | 41577021 | -70640749 | 13/09/16 | 22.63 | 33.26 | 5.67 | 25 |
| Ft. Fisher, North Carolina | 34487519 | -77439831 | 16/03/16 | 22.42 | 31.98 | 5.22 | 28 |

\* Temperatures recorded during the sampling, \*\* temperature calculated from previous weather reports (Reitzel et al., 2013).

**Figure S1.**
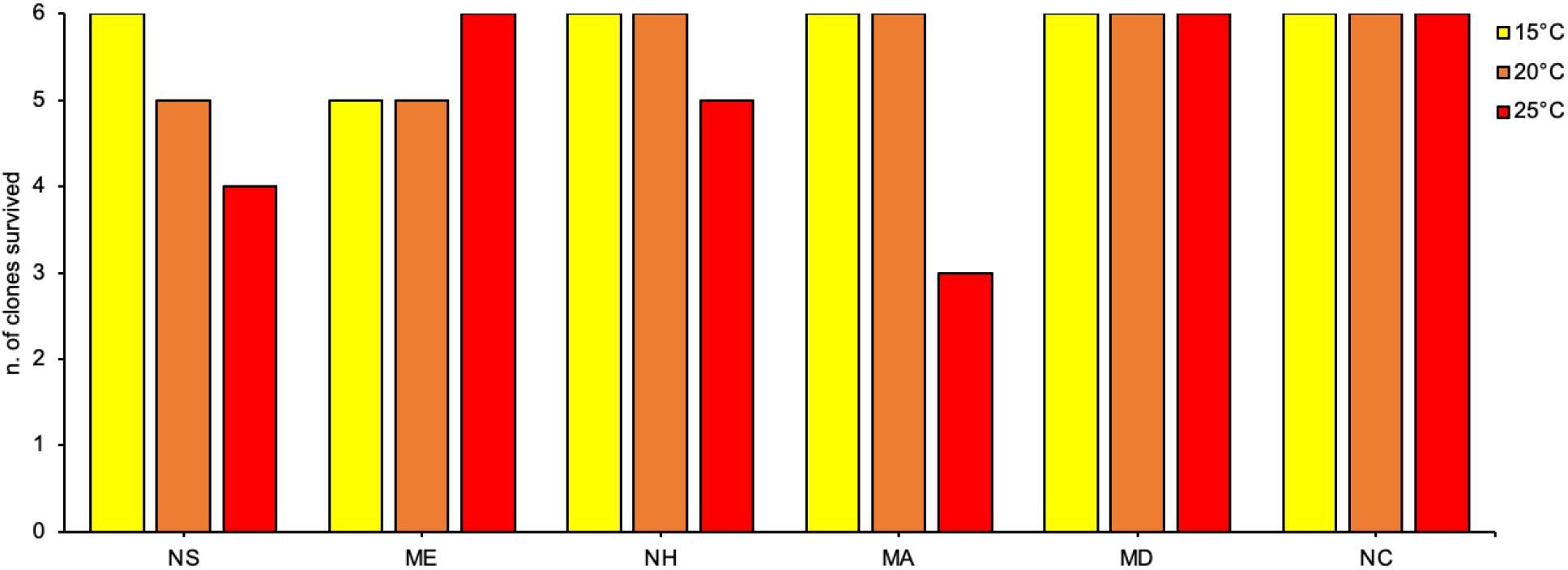
Number of clones for each provenance location that survived at the 3 different temperatures

**Figure S2.**
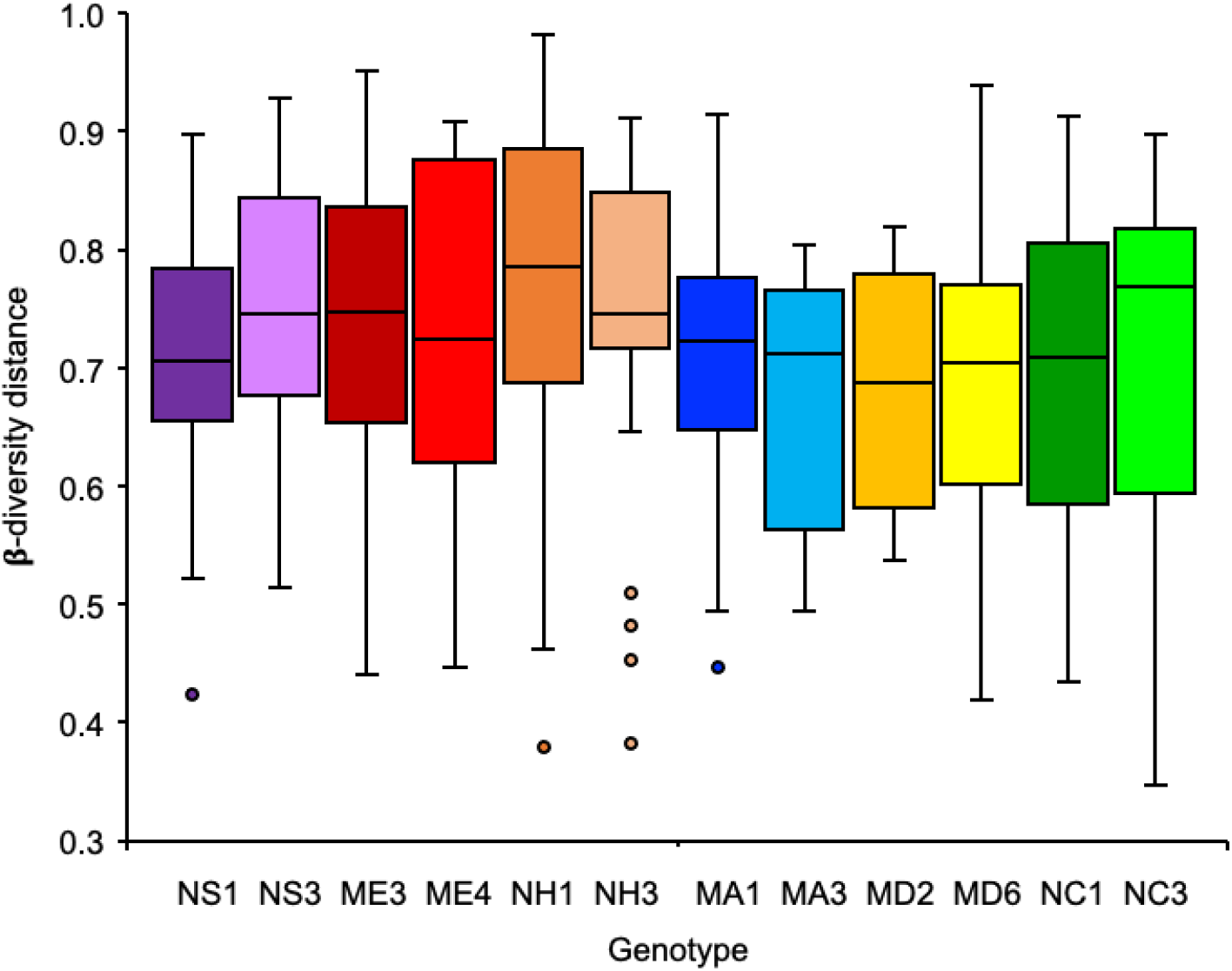
Beta-diversity distance box-plots between different genotypes. (Binary-Pearson metric, sampling depth = 11700), differences were tested through Kruskal-Wallis test (H = 15.54, p = 0.159). NS (Nova Scotia), ME (Maine), NH (New Hampshire), MA (Massachusetts), MD (Maryland), NC (North Carolina), numbers near the location abbreviations indicate the different genotypes.

**Figure S3.**
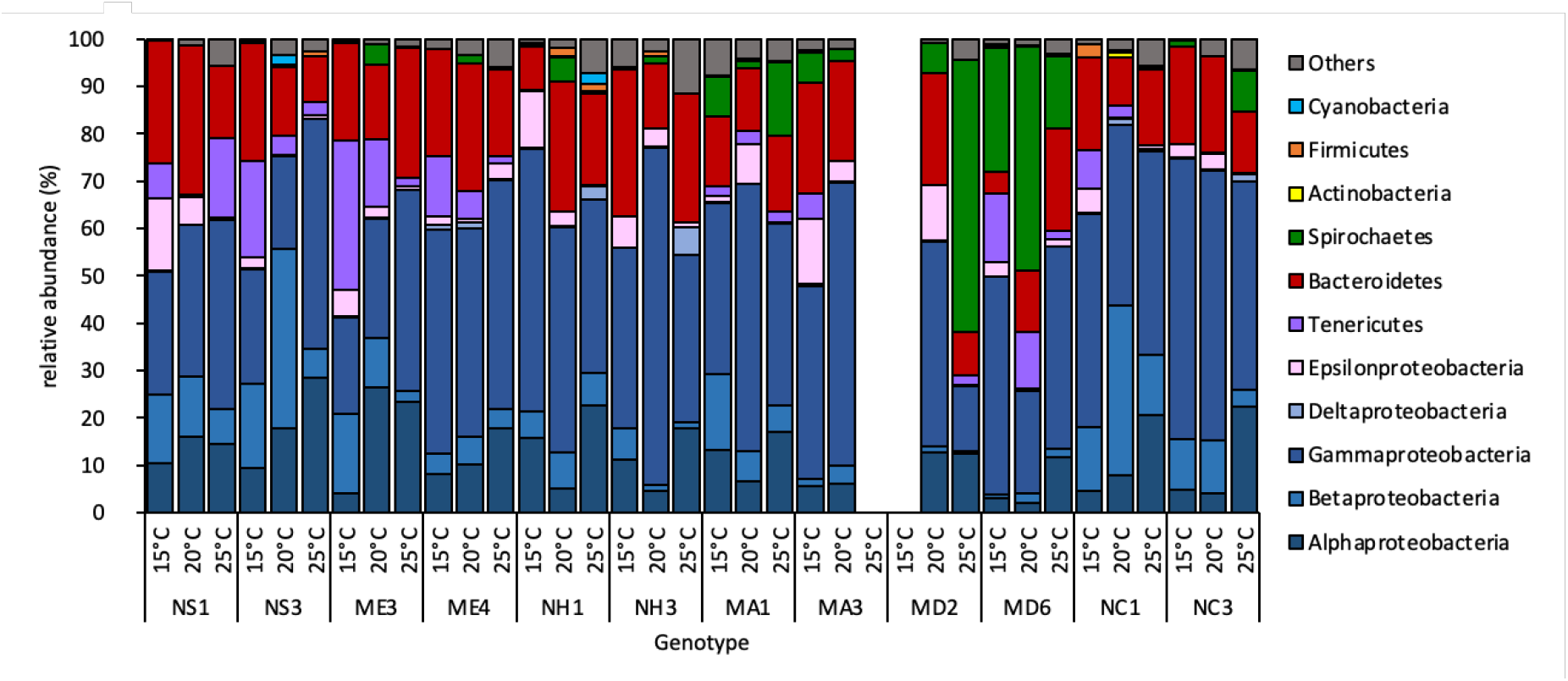
Relative abundance of main bacterial groups among the different genotypes at the three temperatures. NS (Nova Scotia), ME (Maine), NH (New Hampshire), MA (Massachusetts), MD (Maryland), NC (North Carolina), numbers near the location abbreviations indicate the different genotypes.

**Figure S4.**
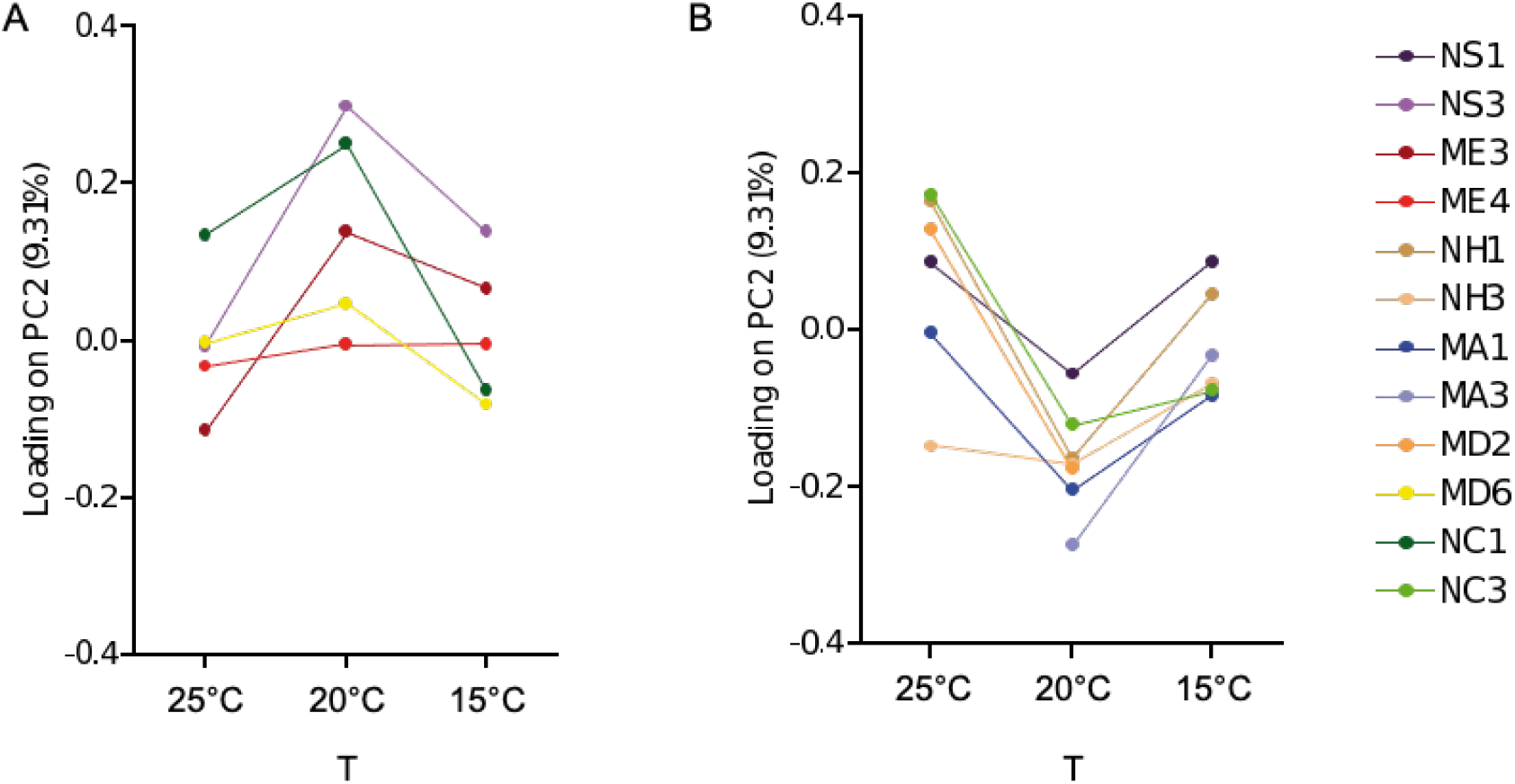
Reaction norms. A and B show the same samples of Fig. 5G divided by similar reaction norms. NS (Nova Scotia), ME (Maine), NH (New Hampshire), MA (Massachusetts), MD (Maryland), NC (North Carolina), numbers near the location abbreviations indicate the different genotypes.

## References

A Hernandez-Agreda, W. L. P. Bongaerts, TD Ainsworth, Hernandez-Agreda, A., Leggat, W., Bongaerts, P., Ainsworth, T. D., & A Hernandez-Agreda, W. L. P. Bongaerts, TD Ainsworth. (2016). The microbial signature provides insight into the mechanistic basis of coral success across reef habitats. MBio, 7(4), e00560–16. https://doi.org/10.1128/mBio.00560-16

Augustin, R., Schröder, K., Rincón, A. P. M., Fraune, S., Anton-Erxleben, F., Herbst, E. M., Wittlieb, J., Schwentner, M., Grötzinger, J., Wassenaar, T. M., & Bosch, T. C. G. (2017). A secreted antibacterial neuropeptide shapes the microbiome of Hydra. Nature Communications, 8(1). https://doi.org/10.1038/s41467-017-00625-1

Baker, A. C. (2003). Flexibility and Specificity in Coral-Algal Symbiosis: Diversity, Ecology, and Biogeography of Symbiodinium. Annual Review of Ecology, Evolution, and Systematics, 34, 661–689. https://doi.org/10.1146/annurev.ecolsys.34.011802.132417

Baldassarre, L., Levy, S., Bar-Shalom, R., Steindler, L., Lotan, T., & Fraune, S. (2021). Contribution of Maternal and Paternal Transmission to Bacterial Colonization in Nematostella vectensis. Frontiers in Microbiology, 12, 2892. https://doi.org/10.3389/fmicb.2021.726795

Baldassarre, L., Ying, H., Reitzel, A., & Fraune, S. (2021). Microbiota mediated plasticity promotes thermal adaptation in Nematostella vectensis [Preprint]. bioRxiv https://doi.org/10.1101/2021.10.18.464790

Bang, C., Dagan, T., Deines, P., Dubilier, N., Duschl, W. J., Fraune, S., Hentschel, U., Hirt, H., Hülter, N., Lachnit, T., Picazo, D., Pita, L., Pogoreutz, C., Rädecker, N., Saad, M. M., Schmitz, R. A., Schulenburg, H., Voolstra, C. R., Weiland-Bräuer, N., … Bosch, T. C. G. (2018). Metaorganisms in extreme environments: Do microbes play a role in organismal adaptation? Zoology, 127, 1–19. https://doi.org/10.1016/j.zool.2018.02.004

Bay, R. A., & Palumbi, S. R. (2015). Rapid Acclimation Ability Mediated by Transcriptome Changes in Reef-Building Corals. https://doi.org/10.1093/gbe/evv085

Bolyen, E., Rideout, J. R., Dillon, M. R., Bokulich, N. A., Abnet, C. C., Al-Ghalith, G. A., Alexander, H., Alm, E. J., Arumugam, M., Asnicar, F., Bai, Y., Bisanz, J. E., Bittinger, K., Brejnrod, A., Brislawn, C. J., Brown, C. T., Callahan, B. J., Caraballo-Rodríguez, A. M., Chase, J., … Caporaso, J. G. (2019). Reproducible, interactive, scalable and extensible microbiome data science using QIIME 2. Nature Biotechnology, 37(8), 852–857. https://doi.org/10.1038/s41587-019-0209-9

Bowerman, K. L., Knowles, S. C. L., Bradley, J. E., Baltrūnaitė, L., Lynch, M. D. J., Jones, K. M., & Hugenholtz, P. (2021). Effects of laboratory domestication on the rodent gut microbiome. ISME Communications, 1(1), 1–14. https://doi.org/10.1038/s43705-021-00053-9

Brener-Raffalli, K., Clerissi, C., Vidal-Dupiol, J., Adjeroud, M., Bonhomme, F., Pratlong, M., Aurelle, D., Mitta, G., & Toulza, E. (2018). Thermal regime and host clade, rather than geography, drive Symbiodinium and bacterial assemblages in the scleractinian coral Pocillopora damicornis sensu lato. Microbiome, 6(1), 39. https://doi.org/10.1186/s40168-018-0423-6

Cahana, I., & Iraqi, F. A. (2020). Impact of host genetics on gut microbiome: Take-home lessons from human and mouse studies. Animal Models and Experimental Medicine, 3(3), 229–236. https://doi.org/10.1002/ame2.12134

Cai, L., Zhou, G., Tong, H., Tian, R. M., Zhang, W., Ding, W., Liu, S., Huang, H., & Qian, P. Y. (2018). Season structures prokaryotic partners but not algal symbionts in subtropical hard corals. Applied Microbiology and Biotechnology, 102(11), 4963–4973. https://doi.org/10.1007/s00253-018-8909-5

Caporaso, J. G., Kuczynski, J., Stombaugh, J., Bittinger, K., Bushman, F. D., Costello, E. K., Fierer, N., Pẽa, A. G., Goodrich, J. K., Gordon, J. I., Huttley, G. A., Kelley, S. T., Knights, D., Koenig, J. E., Ley, R. E., Lozupone, C. A., McDonald, D., Muegge, B. D., Pirrung, M., … Knight, R. (2010). QIIME allows analysis of high-throughput community sequencing data. Nature Methods, 7(5), 335–336. https://doi.org/10.1038/nmeth.f.303

Costello, E. K. EK., Stagaman, K., Dethlefsen, L., Bohannan, BJ. B. J. M., & Relman, D. A. (2012). The application of ecological theory toward an understanding of the human microbiome. Science, 336(6086), 1262. https://doi.org/10.1126/science.1224203

Damjanovic, K., Menéndez, P., Blackall, L. L., & van Oppen, M. J. H. (2020). Early Life Stages of a Common Broadcast Spawning Coral Associate with Specific Bacterial Communities Despite Lack of Internalized Bacteria. Microbial Ecology, 79(3), 706–719. https://doi.org/10.1007/s00248-019-01428-1

Damjanovic, K., Van Oppen, M. J. H., Menéndez, P., & Blackall, L. L. (2019). Experimental Inoculation of Coral Recruits With Marine Bacteria Indicates Scope for Microbiome Manipulation in Acropora tenuis and Platygyra daedalea. Frontiers in Microbiology, 10(JULY). https://doi.org/10.3389/fmicb.2019.01702

Darling, J. A., Kuenzi, A., & Reitzel, A. M. (2009). Human-mediated transport determines the nonnative distribution of the anemone Nematostella vectensis, a dispersal-limited estuarine invertebrate. Marine Ecology Progress Series, 380, 137–146. https://doi.org/10.3354/meps07924

Darling, J. A., Reitzel, A. M., & Finnerty, J. R. (2004). Regional population structure of a widely introduced estuarine invertebrate: Nematostella vectensis Stephenson in New England. Molecular Ecology, 13(10), 2969–2981. https://doi.org/10.1111/j.1365-294X.2004.02313.x

Darling, J. A., Reitzel, A. R., Burton, P. M., Mazza, M. E., Ryan, J. F., Sullivan, J. C., & Finnerty, J. R. (2005). Rising starlet: The starlet sea anemone, Nematostella vectensis. BioEssays, 27(2), 211–221. https://doi.org/10.1002/bies.20181

David, L. A., Maurice, C. F., Carmody, R. N., Gootenberg, D. B., Button, J. E., Wolfe, B. E., Ling, A. V., Devlin, A. S., Varma, Y., Fischbach, M. A., Biddinger, S. B., Dutton, R. J., & Turnbaugh, P. J. (2014). Diet rapidly and reproducibly alters the human gut microbiome. Nature, 505(7484), 559–563. https://doi.org/10.1038/nature12820

Domin, H., Zurita-Gutiérrez, Y. H., Scotti, M., Buttlar, J., Humeida, U. H., & Fraune, S. (2018). Predicted bacterial interactions affect in vivo microbial colonization dynamics in Nematostella. Frontiers in Microbiology, 9(APR). https://doi.org/10.3389/fmicb.2018.00728

Dubé, C. E., Ziegler, M., Mercière, A., Boissin, E., Planes, S., Bourmaud, C. A.-F., & Voolstra, C. R. (2021). Naturally occurring fire coral clones demonstrate a genetic and environmental basis of microbiome composition. Nature Communications, 12(1), 6402. https://doi.org/10.1038/s41467-021-26543-x

Dunphy, C. M., Gouhier, T. C., Chu, N. D., & Vollmer, S. V. (2019). Structure and stability of the coral microbiome in space and time. Scientific Reports, 9(1), 1–13. https://doi.org/10.1038/s41598-019-43268-6

Edwards, S. V. (2020). Genomics of adaptation and acclimation: From field to lab and back. National Science Review, 7(1), 128–128. https://doi.org/10.1093/nsr/nwz173

Faith, J. J., Guruge, J. L., Charbonneau, M., Subramanian, S., Seedorf, H., Goodman, A. L., Clemente, J. C., Knight, R., Heath, A. C., Leibel, R. L., Rosenbaum, M., & Gordon, J. I. (2013). The long-term stability of the human gut microbiota. Science, 341(6141), 1237439–1237439. https://doi.org/10.1126/science.1237439

Franzenburg, S., Fraune, S., Kunzel, S., Baines, J. F., Domazet-Loso, T., Bosch, T. C. G. G., Künzel, S., Baines, J. F., Domazet-Lošo, T., & Bosch, T. C. G. G. (2012). MyD88-deficient Hydra reveal an ancient function of TLR signaling in sensing bacterial colonizers. Proceedings of the National Academy of Sciences, 109(47), 19374–19379. https://doi.org/10.1073/pnas.1213110109

Fraune, S., & Bosch, T. C. G. G. (2007). Long-term maintenance of species-specific bacterial microbiota in the basal metazoan Hydra. Proceedings of the National Academy of Sciences, 104(32), 13146–13151. https://doi.org/10.1073/pnas.0703375104

Friedman, L. E., Gilmore, T. D., & Finnerty, J. R. (2018). Intraspecific variation in oxidative stress tolerance in a model cnidarian: Differences in peroxide sensitivity between and within populations of Nematostella vectensis. PLoS ONE, 13(1), 1–19. https://doi.org/10.1371/journal.pone.0188265

Glasl, B., Smith, C. E., Bourne, D. G., & Webster, N. S. (2019). Disentangling the effect of host-genotype and environment on the microbiome of the coral Acropora tenuis. PeerJ, 7, e6377–e6377. https://doi.org/10.7717/peerj.6377

Goldsmith, D. B., Kellogg, C. A., Morrison, C. L., Gray, M. A., Stone, R. P., Waller, R. G., Brooke, S. D., & Ross, S. W. (2018). Comparison of microbiomes of cold-water corals Primnoa pacifica and Primnoa resedaeformis, with possible link between microbiome composition and host genotype. Scientific Reports, 8(1), 1–15. https://doi.org/10.1038/s41598-018-30901-z

Groussin, M., Mazel, F., Sanders, J. G., Smillie, C. S., Lavergne, S., Thuiller, W., & Alm, E. J. (2017). Unraveling the processes shaping mammalian gut microbiomes over evolutionary time. Nature Communications, 8. https://doi.org/10.1038/ncomms14319

Hand, C., & Uhlinger, K. R. (1992). The culture, sexual and asexual reproduction, and growth of the sea anemone Nematostella vectensis. Biological Bulletin, 182(2), 169–176. https://doi.org/10.2307/1542110

Ibarra-Juarez, L. A., Desgarennes, D., Vázquez-Rosas-Landa, M., Villafan, E., Alonso-Sánchez, A., Ferrera-Rodríguez, O., Moya, A., Carrillo, D., Cruz, L., Carrión, G., López-Buenfil, A., García-Avila, C., Ibarra-Laclette, E., & Lamelas, A. (2018). Impact of Rearing Conditions on the Ambrosia Beetle’s Microbiome. Life, 8(4), 63. https://doi.org/10.3390/life8040063

Kvennefors, E. C. E., Sampayo, E., Ridgway, T., Barnes, A. C., & Hoegh-Guldberg, O. (2010). Bacterial communities of two ubiquitous great barrier reef corals reveals both site-and species-specificity of common bacterial associates. PLoS ONE, 5(4), e10401–e10401. https://doi.org/10.1371/journal.pone.0010401

La Rivière, M., Roumagnac, M., Garrabou, J., & Bally, M. (2013). Transient Shifts in Bacterial Communities Associated with the Temperate Gorgonian Paramuricea clavata in the Northwestern Mediterranean Sea. PLoS ONE, 8(2), e57385. https://doi.org/10.1371/journal.pone.0057385

Lee, S. T. M., Davy, S. K., Tang, S. L., Kench, P. S., & STM Lee, S. D., S. L. Tang, PS Kench. (2016). Mucus sugar content shapes the bacterial community structure in thermally stressed Acropora muricata. 7(MAR), 371–371. https://doi.org/10.3389/fmicb.2016.00371

Leeming, E. R., Johnson, A. J., Spector, T. D., & Le Roy, C. I. (2019). Effect of Diet on the Gut Microbiota: Rethinking Intervention Duration. Nutrients, 11(12), 2862. https://doi.org/10.3390/nu11122862

Li, J., Chen, Q., Long, L. J., Dong, J. D., Yang, J., & Zhang, S. (2014). Bacterial dynamics within the mucus, tissue and skeleton of the coral Porites lutea during different seasons. Scientific Reports, 4. https://doi.org/10.1038/srep07320

Linnenbrink, M., Wang, J., Hardouin, E. A., Künzel, S., Metzler, D., & Baines, J. F. (2013). The role of biogeography in shaping diversity of the intestinal microbiota in house mice. Molecular Ecology, 22(7), 1904–1916. https://doi.org/10.1111/mec.12206

McFall-Ngai, M., Hadfield, M. G., Bosch, T. C. G. G., Carey, H. V., Domazet-Lošo, T., Douglas, A. E., Dubilier, N., Eberl, G., Fukami, T., Gilbert, S. F., Hentschel, U., King, N., Kjelleberg, S., Knoll, A. H., Kremer, N., Mazmanian, S. K., Metcalf, J. L., Nealson, K., Pierce, N. E., … Wernegreen, J. J. (2013). Animals in a bacterial world, a new imperative for the life sciences. Proceedings of the National Academy of Sciences, 110(9), 3229–3236. https://doi.org/10.1073/pnas.1218525110

Morrow, J. L., Frommer, M., Shearman, D. C. A., & Riegler, M. (2015). The Microbiome of Field-Caught and Laboratory-Adapted Australian Tephritid Fruit Fly Species with Different Host Plant Use and Specialisation. Microbial Ecology, 70(2), 498–508. https://doi.org/10.1007/s00248-015-0571-1

Mortzfeld, B. M., Urbanski, S., Reitzel, A. M., Künzel, S., Technau, U., & Fraune, S. (2016). Response of bacterial colonization in Nematostella vectensis to development, environment and biogeography. Environmental Microbiology, 18(6), 1764–1781. https://doi.org/10.1111/1462-2920.12926

Oyserman, B. O., Cordovez, V., Flores, S. S., Leite, M. F. A., Nijveen, H., Medema, M. H., & Raaijmakers, J. M. (2021). Extracting the GEMs: Genotype, Environment, and Microbiome Interactions Shaping Host Phenotypes. Frontiers in Microbiology, 11. https://www.frontiersin.org/article/10.3389/fmicb.2020.574053

Pearson, C. V. M., Rogers, A. D., & Sheader, M. (2002). The genetic structure of the rare lagoonal sea anemone, Nematostella vectensis Stephenson (Cnidaria; Anthozoa) in the United Kingdom based on RAPD analysis. Molecular Ecology, 11(11), 2285–2293. https://doi.org/10.1046/j.1365-294X.2002.01621.x

Peixoto, R. S., Rosado, P. M., Leite, D. C. de A., Rosado, A. S., & Bourne, D. G. (2017). Beneficial microorganisms for corals (BMC) Proposed mechanisms for coral health and resilience. Frontiers in Microbiology, 8(MAR), 341. https://doi.org/10.3389/fmicb.2017.00341

Putnam, N. H., Srivastava, M., Hellsten, U., Dirks, B., Chapman, J., Salamov, A., Terry, A., Shapiro, H., Lindquist, E., Kapitonov, V. V., Jurka, J., Genikhovich, G., Grigoriev, I. V., Lucas, S. M., Steele, R. E., Finnerty, J. R., Technau, U., Martindale, M. Q., & Rokhsar, D. S. (2007). Sea anemone genome reveals ancestral eumetazoan gene repertoire and genomic organization. Science, 317(5834), 86–94. https://doi.org/10.1126/science.1139158

Rausch, P., Basic, M., Batra, A., Bischoff, S. C., Blaut, M., Clavel, T., Gläsner, J., Gopalakrishnan, S., Grassl, G. A., Günther, C., Haller, D., Hirose, M., Ibrahim, S., Loh, G., Mattner, J., Nagel, S., Pabst, O., Schmidt, F., Siegmund, B., … Baines, J. F. (2016). Analysis of factors contributing to variation in the C57BL/6J fecal microbiota across German animal facilities. International Journal of Medical Microbiology, 306(5), 343–355. https://doi.org/10.1016/j.ijmm.2016.03.004

Reitzel, A. M., Burton, P. M., Krone, C., & Finnerty, J. R. (2007). Comparison of developmental trajectories in the starlet sea anemone Nematostella vectensis: Embryogenesis, regeneration, and two forms of asexual fission. Invertebrate Biology, 126(2), 99–112. https://doi.org/10.1111/j.1744-7410.2007.00081.x

Reitzel, A. M., Chu, T., Edquist, S., Genovese, C., Church, C., Tarrant, A. M., & Finnerty, J. R. (2013). Physiological and developmental responses to temperature by the sea anemone Nematostella vectensis. Marine Ecology Progress Series, 484(Somero 2012), 115–130. https://doi.org/10.3354/meps10281

Reitzel, A. M., Darling, J. A., Sullivan, J. C., & Finnerty, J. R. (2008). Global population genetic structure of the starlet anemone Nematostella vectensis: Multiple introductions and implications for conservation policy. Biological Invasions, 10(8), 1197–1213. https://doi.org/10.1007/s10530-007-9196-8

Reitzel, A. M., Herrera, S., Layden, M. J., Martindale, M. Q., & Shank, T. M. (2013). Going where traditional markers have not gone before: Utility of and promise for RAD sequencing in marine invertebrate phylogeography and population genomics. Molecular Ecology, 22(11), 2953–2970. https://doi.org/10.1111/mec.12228

Reitzel, A. M., Sullivan, J. C., & Finnerty, J. R. (2010). Discovering SNPs in protein coding regions with stellasnp: Illustrating the characterization and geographic distribution of polymorphisms in the estuarine anemone nematostella vectensis. Estuaries and Coasts, 33(4), 930–943. https://doi.org/10.1007/s12237-009-9231-3

Reshef, L., Koren, O., Loya, Y., Zilber-Rosenberg, I., & Rosenberg, E. (2006). The Coral Probiotic Hypothesis. Environmental Microbiology, 8(12), 2068–2073. https://doi.org/10.1111/j.1462-2920.2006.01148.x

Ritchie, K. K. B. (2006). Regulation of microbial populations by coral surface mucus and mucus-associated bacteria. 322, 1–14. https://doi.org/10.3354/meps322001

Rosado, P. M., Leite, D. C. A., Duarte, G. A. S., Chaloub, R. M., Jospin, G., Nunes da Rocha, U., P. Saraiva, J., Dini-Andreote, F., Eisen, J. A., Bourne, D. G., & Peixoto, R. S. (2019). Marine probiotics: Increasing coral resistance to bleaching through microbiome manipulation. ISME Journal, 13(4), 921–936. https://doi.org/10.1038/s41396-018-0323-6

Rubio-Portillo, E., Santos, F., Martínez-García, M., de los Ríos, A., Ascaso, C., Souza-Egipsy, V., Ramos-Esplá, A. A., & Anton, J. (2016). Structure and temporal dynamics of the bacterial communities associated to microhabitats of the coral Oculina patagonica. Environmental Microbiology, 18(12), 4564–4578. https://doi.org/10.1111/1462-2920.13548

Segata, N., Izard, J., Waldron, L., Gevers, D., Miropolsky, L., Garrett, W. S., & Huttenhower, C. (2011). Metagenomic biomarker discovery and explanation. Genome Biology, 12(6), R60–R60. https://doi.org/10.1186/gb-2011-12-6-r60

Sehnal, L., Brammer-Robbins, E., Wormington, A. M., Blaha, L., Bisesi, J., Larkin, I., Martyniuk, C. J., Simonin, M., & Adamovsky, O. (2021). Microbiome Composition and Function in Aquatic Vertebrates: Small Organisms Making Big Impacts on Aquatic Animal Health. Frontiers in Microbiology, 12, 358. https://doi.org/10.3389/fmicb.2021.567408

Sharp, K. H., Pratte, Z. A., Kerwin, A. H., Rotjan, R. D., & Stewart, F. J. (2017). Season, but not symbiont state, drives microbiome structure in the temperate coral Astrangia poculata. Microbiome, 5(1), 120–120. https://doi.org/10.1186/s40168-017-0329-8

Sheader, M., Suwailem, A. M., & Rowe, G. A. (1997). The anemone, Nematostella vectensis, in Britain: Considerations for conservation management. Aquatic Conservation: Marine and Freshwater Ecosystems, 7(1), 13–25. https://doi.org/10.1002/(SICI)1099-0755(199703)7:1<13::AID-AQC210>3.0.CO;2-Y

Smith, K. J., & Able, K. W. (1994). Salt-marsh tide pools as winter refuges for the mummichog, Fundulus heteroclitus, in New Jersey. Estuaries, 17(1), 226–234. https://doi.org/10.2307/1352572

Staubach, F., Baines, J. F., Künzel, S., Bik, E. M., & Petrov, D. A. (2013). Host Species and Environmental Effects on Bacterial Communities Associated with Drosophila in the Laboratory and in the Natural Environment. PLOS ONE, 8(8), e70749. https://doi.org/10.1371/journal.pone.0070749

Terraneo, T. I., Fusi, M., Hume, B. C. C., Arrigoni, R., Voolstra, C. R., Benzoni, F., Forsman, Z. H., & Berumen, M. L. (2019). Environmental latitudinal gradients and host-specificity shape Symbiodiniaceae distribution in Red Sea Porites corals. Journal of Biogeography, 46(10), 2323–2335. https://doi.org/10.1111/jbi.13672

van Oppen, M. J. H., Bongaerts, P., Frade, P., Peplow, L. M., Boyd, S. E., Nim, H. T., & Bay, L. K. (2018). Adaptation to reef habitats through selection on the coral animal and its associated microbiome. Molecular Ecology, 27(14), 2956–2971. https://doi.org/10.1111/mec.14763

Vijayan, N., Lema, K. A., Nedved, B. T., & Hadfield, M. G. (2019). Microbiomes of the polychaete Hydroides elegans (Polychaeta: Serpulidae) across its life-history stages. Marine Biology, 166(2). https://doi.org/10.1007/s00227-019-3465-9

Voolstra, C. R., & Ziegler, M. (2020). Adapting with Microbial Help Microbiome Flexibility Facilitates Rapid Responses to Environmental Change. BioEssays, 42(7), 2000004. https://doi.org/10.1002/bies.202000004

Zarrinpar, A., Chaix, A., Yooseph, S., & Panda, S. (2014). Diet and feeding pattern affect the diurnal dynamics of the gut microbiome. Cell Metabolism, 20(6), 1006–1017. https://doi.org/10.1016/j.cmet.2014.11.008

Ziegler, M., Grupstra, C. G. B., Barreto, M. M., Eaton, M., BaOmar, J., Zubier, K., Al-Sofyani, A., Turki, A. J., Ormond, R., & Voolstra, C. R. (2019). Coral bacterial community structure responds to environmental change in a host-specific manner. Nature Communications, 10(1), 3092. https://doi.org/10.1038/s41467-019-10969-5

Ziegler, M., Seneca, F. O., Yum, L. K., Palumbi, S. R., & Voolstra, C. R. (2017). Bacterial community dynamics are linked to patterns of coral heat tolerance. Nat. Commun., 8, 14213–14213. https://doi.org/10.1038/ncomms14213

Zilber-Rosenberg, I., & Rosenberg, E. (2008). Role of microorganisms in the evolution of animals and plants: The hologenome theory of evolution. FEMS Microbiology Reviews, 32(5), 723–735. https://doi.org/10.1111/j.1574-6976.2008.00123.x

